# Stromal cell subsets modulate T-cell infiltration in early breast cancer

**DOI:** 10.1101/2025.10.20.683407

**Authors:** Julia Chen, Hanyun Zhang, Travis Ruan, Sunny Z Wu, Iveta Slapetova, Ewan Millar, Peter H Graham, Jodi Lynch, Lois H Browne, Elgene Lim, Alexander Swarbrick

## Abstract

Recent studies of the tumour microenvironment have elucidated the heterogeneity of stromal cells, with increasing evidence suggesting that stromal subsets play an important role in regulating anti-tumour immunity in breast cancer. However, the functional diversity of these cells within the tumour immune microenvironment and how they interact with immune cells in a spatial and clinical context remain poorly understood. We performed multiplex immunofluorescence on tumour microarrays from two cohorts consisting of 591 breast cancer patients to assess the abundance and spatial co-localisation of stromal and immune subsets and their correlation with clinicopathological features and patient outcomes. We found that stromal cells were spatially distinct. A perivascular-like subset that was disseminated throughout the stroma rather than restricted to vessel-adjacent regions was enriched in an immune cold environment and associated with T-cell exclusion. An inflammatory-like cancer-associated fibroblast subset was associated with segregation of T cells from cancer cells. Our findings highlight the differential impact of stromal subsets on immune infiltration and activation within the breast cancer tumour microenvironment with potential implications for patient outcomes.

**Significance:** This study characterised spatially-defined interactions between stromal and immune subsets in large clinical cohorts of early breast cancer. Findings from this study will contribute to gaps in current knowledge in how diverse stromal cell subsets, particularly novel subsets, interact with immune cells in a clinically-relevant context.

## Introduction

Breast cancer is the most commonly diagnosed cancer in women and despite advancements in treatment over the last few decades, it remains the leading cause of cancer-related death among females worldwide (1). Recent studies into the tumour microenvironment (TME) have revealed complex functional roles of stromal cells in tumorigenesis, progression and suppression (2). Although there is increasing evidence that stromal cells affect tumour immunity, the diversity of stromal cell subsets and their interactions with immune cells in a clinically relevant context remain poorly understood.

Recent advances in technologies such as single-cell RNA-sequencing (scRNA-seq) have allowed the characterisation of tumours in tremendous resolution and results highlight the heterogeneity of stromal and immune cells within the TME. Different cancer-associated fibroblast (CAF) subpopulations or states with discrete phenotypic, transcriptional and functional properties have been described in various cancer types (3–7). In breast cancer, our previous scRNA-seq study has revealed four distinct populations of stromal cells: myofibroblast-like CAF (myCAF), inflammatory-like CAF (iCAF) and two perivascular-like cells (PVLs) in differentiated and immature states (8, 9). Each of these subpopulations displayed distinct morphologies, spatial relationships and functional properties. Similarly, scRNA-seq of immune cells showed that a significant heterogeneity exists in the infiltrating T cell population (10). The effectiveness of CD8^+^ T cells is regulated by immune checkpoint molecules including PD-1 and its ligand PD-L1 and these are the targets of most currently available immunotherapies.

One of the major limitations of single-cell technologies is the lack of information on the structural organisation of cells within a tumour, which has increasingly been recognised as a factor that contributes to the heterogeneity, variable tumour behaviours and differential response to treatments observed in most cancers (11). Spatial profiling has hence been explored to provide a more comprehensive view of the intricate cellular makeup of solid tumours, enabling examination of different cell types, their locations and interactions within different layers of spatial organisation. Integration of this with clinical data from large cohorts enables the exploration of spatially defined TME components as predictors of treatment response and patient outcomes, with the potential to discover novel biomarkers and therapeutic targets, thereby improving the precision of cancer diagnosis and management (12).

Here we performed multiplex immunofluorescence (mIF) on tumour microarrays (TMAs) from two independent primary breast cancer cohorts using a panel of stromal and immune markers defined using our scRNA-seq results. The abundance and spatial co-localisation of annotated stromal and immune subsets were linked with clinicopathological data to assess their clinical significance.

## MATERIALS AND METHODS

### Clinical cohorts

This study consists of TMA cores from 591 breast cancer patients with luminal and triple-negative breast cancer (TNBC) subtypes. All tumours were sampled pre-treatment. Luminal patients (n=369) were recruited as part of the Breast Boost trial where patients received wide local excision with whole breast irradiation with randomisation to cavity boost or no boost (13). The TNBC cases (n=222) were identified by a review of the Oncology database at St George Hospital between 2004 and 2019 (14). Tumours were classified into molecular subtypes according to the following definitions: luminal A: ER^+^ ≥1%, PR^+^>20%, HER2^-^, Ki67 < 14%; luminal B: ER^+^, PR^+^≤ 20% and/or HER2^+^ and/or Ki67 ≥14%; TNBC: ER/PR^-^ (<1%), HER2^-^.

TMA slides were constructed with 3×1mm cores sampled from the periphery of tumour blocks marked by a breast pathologist on a H&E slide. Paraffin sections were cut at 4μm onto Superfrost glass slides (ThermoFisher Scientific, Waltham, MA, USA) and stained using H&E using a Leica Bond Rx automated immunostainer machine (Leica Biosystems, Nussloch, Germany).

### Multiplexed immunofluorescence

Opal 9^TM^ was used for mIF imaging (Akoya Biosicences, Marlborough, MA, USA), with primary antibody conditions optimised for DAB (3,3’-diaminobenzidine) staining applied to both monoplex and multiplex optimisations. Each biomarker antibody was paired with a specific Opal fluorophore based on the co-expression patterns of biomarkers in the tissue and their anticipated protein expression levels (**Supplementary table 1**).

### Image analysis

Slides were scanned using the Vectra Polaris 3.0 (Akoya Biosciences) using 40 × magnification. Cell type identification and classification were performed in accordance with the method described by Wang et al (14). Briefly, TMA images were analysed using QuPath v0.2.3 (RRID:SCR_018257, https://qupath.github.io). All TMA files and their corresponding TMA maps were imported using the TMA module. Tissue core identification numbers generated by the TMA de-arrayer were verified before analysis. Tissue regions were segmented into tumour and stromal compartments using a pixel-based machine learning classifier trained using PanCK staining. Cell segmentation was based on DAPI nuclear staining and implemented using the inbuilt cell detection algorithm. Biomarker phenotyping was conducted through object classification using a random forest algorithm applied to mean nuclear intensity measurements to minimise staining leakage from nearby cells. The final combined classifier was applied to all TMA cores. Cell types were defined by expression or co-expression of markers, including epithelial (PanCK^+^), endothelial (CD31^+^), myCAF (CD140b^+^αSMA^+^), iCAF (CD140b^+^aSMA^-^CD146^-^), differentiated-PVLs (dPVLs) (CD146^+^THY1^-^), immature-PVL (imPVLs) (CD146^+^THY1^+^), PD1^+^ CD8 T cells (PD1^+^CD8^+^), and other CD8 T cells (PD1^-^CD8^+^) (**Supplementary table 2**).

TMA core measurements were linked with corresponding clinical data using patient ID. For patients with more than one core, the median values of these cores were used for analysis. The abundance of cell types was measured for each core as the percentage of cells per total number of detections. Cell proportions were computed within the stromal and tumour regions separately. To account for differences in locations, the proportion of immune cells were taken from the entire core whereas for stromal cells, proportions were taken from stromal regions only to reduce the impact of misclassified stromal cells in tumour regions.

### Spatial analysis

To investigate the spatial proximity between cell types, we measured the proportion of cell type A with nearby cell type B within a radius of 30-100 µm, with an interval of 10 µm. Cell type A with at least one cell type B within 30 µm were determined as B-adjacent cells. Type A cells without type B cells within 100 µm were defined as B-distal cells. We also measured the minimal distance between a pair of cell types using the *calculate_minimum_distances_between_celltypes* function from the SPIAT package (15). The patient-level metrics were summarised as the median across TMA cores.

### Immune phenotypes

To characterise patterns of immune distribution, we classified cores into three categories, immune cold, immune intermixing, and immune segregated. Immune cold was determined by a CD8^+^ T cell infiltration (sum of PD1^+^CD8^+^ and PD1^-^CD8^+^ proportions) below the median. We then examined the co-localisation of CD8^+^ T and epithelial cells using the area under curve (AUC) of cross K function (15). A positive AUC score of cross K function indicates co-localisation of these cells while a negative AUC score indicates a separation of CD8^+^ T from epithelial cells. We classified cores with CD8^+^ T cell proportion over the median and a negative cross K AUC as immune segregated. These cores displayed immune aggregates forming outside the epithelial nest. An immune intermixing group was defined with CD8^+^ T proportion over the median and a positive cross K AUC.

### Association of spatial features with clinical variables

We compared the enrichment of stromal subset-related features in groups of patients stratified by clinical characteristics. Features include fibroblast proportions in the stromal region, minimal distance between pairs of cells, proportion of cell type A with cell type B present within a 30μm neighbourhood, and the normalised mixing score defined by the interactions between cell type A and B divided by the sum of interactions between A-A and B-B (15) (**Supplementary table 3**). The Wilcoxon rank sum test was performed using the *compare_means* function from ggpubr (0.6.0, RRID:SCR_021139) for each feature to determine its difference between groups classified by clinical characteristics. P-value was corrected using the Benjamini-Hochberg (BH) method.

### Statistical Analysis

Statistical analysis was completed using R 4.4.2. Univariate and multivariate survival analyses were conducted using COX proportional hazards modelling, with a p-value <0.05 considered significant. Clinical variables with known prognostic values, including age, lymph node metastasis, grade, tumour size, and molecular subtypes (luminal A vs. luminal B) were included in the multivariate survival analyses. Survival predictions were represented using Kaplan-Meier (KM) plots (survival 3.7.0, RRID:SCR_021137). Association between specific stromal and immune phenotypes and clinicopathological features were examined using Krustal-Wallis tests and Pearson correlations (stats 4.4.2, RRID:SCR_025678). Statistical significance for comparing patient groups was determined by a p-value <0.05 using the Wilcoxon rank sum test.

### Xenium data generation

Formalin-fixed Paraffin-embedded (FFPE) blocks from three TNBC tumours were sectioned at 5 µm and placed onto Xenium slides following the Xenium *In Situ* FFPE Tissue Preparation Guide (10x Genomics, CG000578). Slides were then deparaffinised and de-crosslinked (10x Genomics, CG000580). A predesigned and add-on gene panel of 5101 genes (100 custom add-on genes additional to the default 10X 5k panel; **Supplementary Table 4**) was added to the tissue following instructions in Xenium Prime (5K) *In situ* Gene Expression (10x Genomics, CG000760). Cell segmentation reagents were applied to assist segmentation of nuclei and cell body (10x Genomics, CG000760). Xenium slides were processed on the Xenium Analyser (serial number XETG00531, software version 3.4.1.0, software analysis version xenium-3.3.0.1) for imaging and analysis (Decoding and Imaging user guide, 10x Genomics, CG000584).

### Xenium cell type annotation and spatial analysis

Raw Xenium data were converted into AnnData objects using anndata (v5.2.1) (16). Cells with fewer than 10 detected transcripts or fewer than 1 expressed gene were excluded. Cell annotation was performed based on a combined positivity of genes corresponding to the protein markers in the mIF panel (**Supplementary Table 5**). To identify epithelial cells, *EPCAM* was used as a surrogate of keratin genes encoding PanCK. Adjacent and disseminated PVLs were identified as PVLs (dPVL or imPVL) localised within 30 µm or beyond 100 µm to the nearest endothelial cell, respectively. Distance from PD1^+^CD8^+^ and PD1^-^CD8^+^ T cells to the nearest adjacent or disseminated PVLs was computed using *var_by_distance* function in Squidpy (RRID:SCR_026157) to assess the spatial proximity of T cells to PVL phenotypes (17).

### Differential gene expression and pathway enrichment

The differential gene expression (DEG) analysis was performed using DESeq2 (1.46.0, RRID:SCR_000154) (18). To reduce the false discovery of DEGs arising from transcript contamination from nearby cells, we constrained DEGs to genes expressed by at least 5% of PVLs in a breast cancer scRNA-seq dataset (8). Gene Ontology (GO) enrichment analysis was computed separately for upregulated and downregulated genes comparing between adjacent and disseminated PVLs. Genes were first filtered by significance (absolute log_2_ fold change > 0.5 and adjusted P-value < 0.05), then the top 100 genes from each subset (ranked by log_2_ fold change) were analysed with clusterProfiler (4.14.6, RRID:SCR_016884) (19). Enrichment was performed for Biological Process terms with BH adjustment.

### Ethics

The Breast Boost study was approved by the Human Research Ethics Committee of St George Hospital, Sydney, Australia (ref No: 96/84), and written informed consent was obtained from all participants. Ethics approval for the TNBC cohort was granted by the South Eastern Sydney Local Health District Human Research Ethics Committee at the Prince of Wales Hospital, Sydney (Boost: HREC 96/16 and TNBC: HREC 2018/ETH00138), who granted a waiver of consent to analyse the tissue blocks. The TNBC samples profiled using Xenium Prime in this study were collected following protocols x13-0133, x16-018 and x19-0496. Ethical approval for this study was acquired by the Sydney Local Health Districts Ethics committee (Royal Prince Alfred Hospital zone) and St Vincent’s hospital Ethics Committee. Consent for the use of samples in this study was obtained from all patients prior to collection of tissue, and data were de-identified as per approved protocol. All studies were conducted in accordance with the Declaration of Helsinki.

### Data availability

Raw mIF images and Xenium Prime data are available at https://doi.org/10.6019/S-BSST3074. Processed data can be obtained from https://doi.org/10.5281/zenodo.20199307. Other data generated in this study are available from the corresponding author upon reasonable request.

### Code availability

Code related to the analyses in this study can be found on GitHub at https://github.com/Swarbricklab-code/Stromal_Tcell_Infiltration_mIF

## RESULTS

### Patient demographics

A total of 1,356 TMA cores were processed, with a median of 3 cores per patient (range 1-6). 868 TMA cores were from luminal tumours and 488 cores from TNBC tumours (**Figure 1A**). In the luminal cohort, the majority of tumours were luminal A (n=275, 75%) and invasive ductal carcinoma (n = 317, 85.9%). Other histology subtypes include lobular carcinoma (n=34, 9.2%), micropapillary (n = 7, 1.9%), mucinous (n=10, 2.7%) and tubule-lobular (n = 1, 0.3%). 50% (n=184) of patients in this cohort received adjuvant endocrine therapy, 16% (n=59) received chemotherapy and 10% (n=37) received both. The median length of follow-up for this cohort is 16 years for survival. Analysis of clinicopathological features revealed that age, nodal status and molecular subtypes (luminal A vs. luminal B) were independent predictors of patient survival when adjusted for other clinical factors, consistent with known results from literature (**Supplementary table 6**). The interventional endpoint of this trial, boost vs no boost had no impact on patient outcome (13). For the TNBC cohort, the majority of patients had ductal histology (n=204, 91.9%) and node-negative disease (n=139, 62.6%). 70.3% of patients received adjuvant chemotherapy, including both anthracycline-based and anthracycline-free regimens. The median length of follow-up was 4.5 years for survival. Older age, node positivity, no chemotherapy and large tumour size were associated with poorer survival whereas a high tumour-infiltrating lymphocyte (TIL) score was associated with better survival in the univariate analyses. Multivariate analysis showed a significant correlation between age and nodal status and overall survival (OS) after adjusting for other significant clinical characteristics as confounders (**Supplementary table 7**).

**Figure 1.**
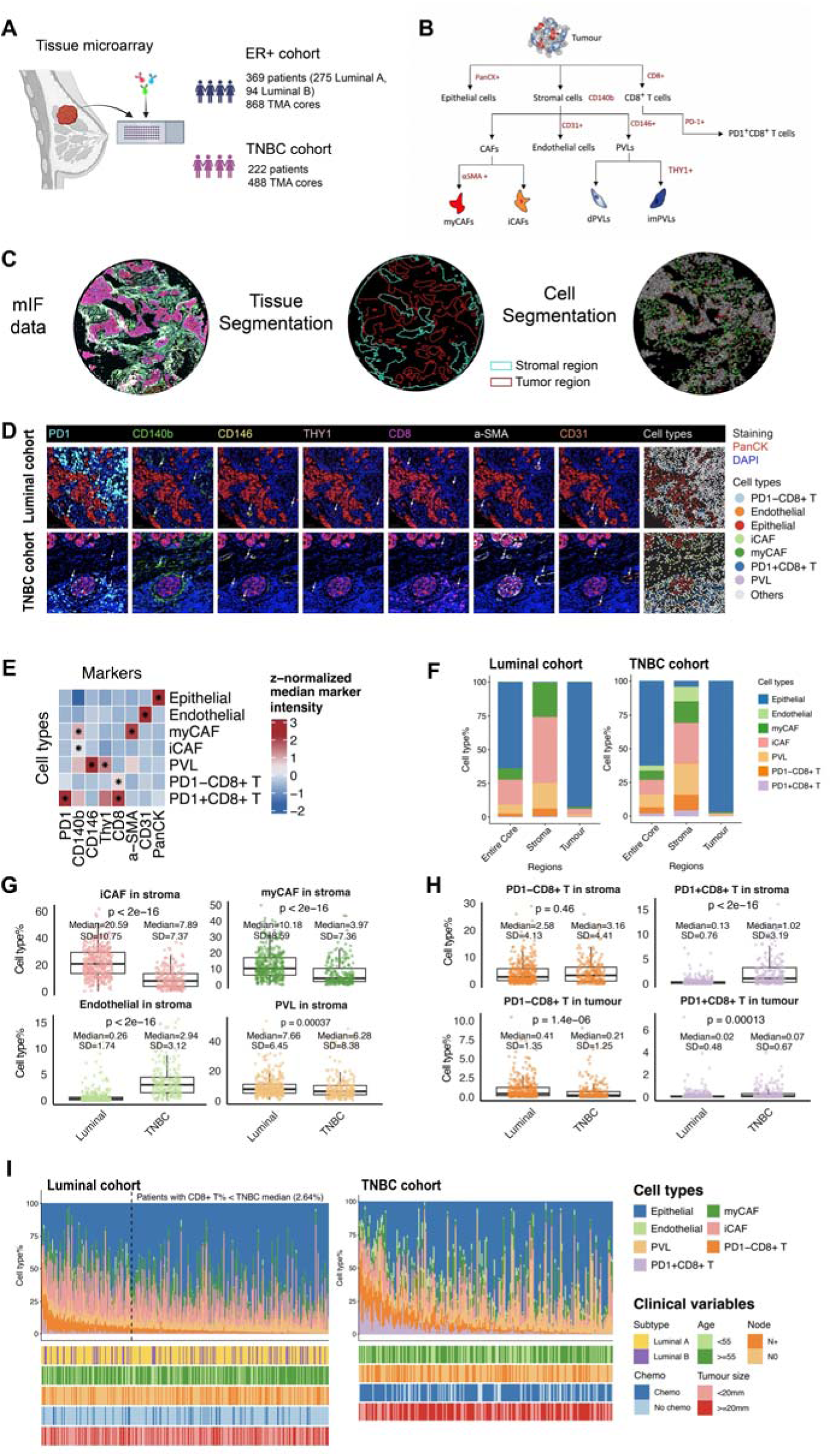
Cohort description and cell type annotations. **A.** Cohort overview (Created in BioRender. Zhang, H. (2026) https://BioRender.com/y2inm3s, RRID:SCR_018361). **B.** Cell types of interest defined by the expression of markers. **C.** mIF images were processed with tissue segmentation and cell segmentation to map the distribution of individual cells in stromal and tumour regions. **D.** mIF images showing marker staining and cell type annotation. **E.** Heatmap showing z-score-normalised median intensities of markers across cell types. Stars indicate the key markers of cell types used for classification. **F.** Median percentages of cell types within cells with assigned classes across the entire TMA core, stromal and tumour regions. **G.** Proportions of iCAFs, myCAFs, endothelial cells, and PVLs within all detected cells in the stromal region in luminal and TNBC cohorts. **H.** Proportions of PD1^-^CD8^+^ T cells and PD1^+^CD8^+^ T cells within all detected cells in the stromal and tumour regions in luminal and TNBC cohorts. **I.** Compositions of classified cell types in the entire TMA cores per patient, with unclassifiable cells excluded. In the luminal cohort, the vertical dotted line separates patients with CD8 T% above and below the median CD8 T% of the TNBC cohort. Colour bars below the bar plot denote the clinical variables per patient.

### Distinct stromal and immune landscapes in luminal and triple-negative breast cancer

mIF data were processed using tissue and cell segmentation to locate individual cells in the tumour or stromal regions. Cells were classified into one of the eight types based on the expression of eight markers, including epithelial (PanCK^+^), endothelial (CD31^+^), myCAF (CD140b^+^αSMA^+^), iCAF (CD140b^+^α SMA^-^CD146^-^), dPVLs (CD146^+^THY1^-^), imPVLs (CD146^+^THY1^+^), PD1^+^ CD8 T cells (PD1^+^CD8^+^), and other CD8 T cells (PD1^-^CD8^+^) (**Figure 1B-D**). The marker intensity was consistent with the cell type definitions (**Figure 1E**).

Cell proportions were quantified as the number of cells normalised by the total number of detected cells within the entire TMA core, as well as within tumour and stromal regions separately. Compositions of classified cells were showed in **Figure 1F**. Proportions per patient were summarised as the median across cores. Due to the sparsity of imPVL, we merged dPVL and imPVL into a general PVL group for downstream analyses. TNBC tumours were enriched in endothelial cells compared to luminal tumours, while the proportion of iCAF, myCAF, and PVL were significantly elevated in luminal tumours (p<0.05, **Figure 1G**).

Within the stromal compartment, iCAF represented the most predominant cell type in both luminal and TNBC cases (median 20.59% in luminal and 7.89% in TNBC, **Figure 1G**). The least abundant cell type is imPVL, accounting for 0.036% in Luminal and 0.041% in TNBC. PVLs account for 7.66% (SD 6.45%) and 6.28% (SD 8.38%) of total detected cells in luminal and TNBC respectively, with a standard deviation comparable to that of iCAF and myCAF (iCAF, SD 10.75% in luminal, 7.37% in TNBC; myCAF, SD 8.59% in luminal, 7.36% in TNBC, **Figure 1G**). In contrast, endothelial cells showed relatively stable abundance across patients (SD 1.73% in luminal, 3.12% in TNBC), despite their presumed spatial association with PVL cells (**Figure 1G**).

In both stromal and tumour regions, TNBC tumours harboured a higher abundance of PD1^+^CD8^+^ T cells than luminal cases (p<0.05, **Figure 1H**), aligning with the notion that TNBC is a more immune-inflamed subtype and can benefit from immune checkpoint therapy (20, 21). Surprisingly, luminal cases showed a higher level of PD1^-^CD8^+^ T in the tumour region than TNBC (p<0.001, **Figure 1H**). Despite luminal tumours containing a lower proportion of CD8^+^ T in general than TNBC (1.58% vs. 2.64%), 116 out of 369 (31.4%) luminal tumours showed a CD8^+^ T cell percentage higher than the median of TNBC tumours (**Figure 1I**), highlighting heterogeneity in immune infiltration within luminal tumours. Additionally, CD8^+^ T cell percentage in luminal tumours increased with younger age at diagnosis (p=0.025, **Supplementary figure 1A**), no lymph node metastasis (p=0.013, **Supplementary figure 1B**), and low grade compared to median grade (p=0.045, **Supplementary figure 1C**). No significant difference in CD8^+^ T cell percentage was observed between luminal A and luminal B subtypes (**Supplementary figure 1D**), or between small (<20mm) and large (>=20mm) tumours (**Supplementary figure 1E**).

We calculated the ratio of PD1^+^CD8^+^ to total CD8^+^ T cells to quantify the prevalence of CD8^+^ T cells expressing inhibitory receptors. This ratio varied across patients, ranging from 0 to 0.55 in luminal (median = 0.05), and 0 to 1 (median = 0.26) in TNBC. Patients with higher overall CD8^+^ T cell proportions tend to have a higher PD1^+^CD8^+^ ratio (luminal, Pearson’s R = 0.15, p = 0.005; TNBC, Pearson’s R = 0.22, p = 0.00086), suggesting that tumours more permissive of T cell infiltration also exhibit increased frequency of inhibitory T cell phenotypes.

### Spatially distinct distribution of stromal subsets and correlation with T cell infiltration

The three stromal subsets exhibited distinct spatial distributions. myCAFs were most closely associated with cancer cells, as indicated by their shortest median distance to tumour cells and their highest representation in cancer-adjacent compartments (**Figure 2A**). In contrast, iCAFs and PVL cells were predominantly located at greater distances from tumour regions.

**Figure 2.**
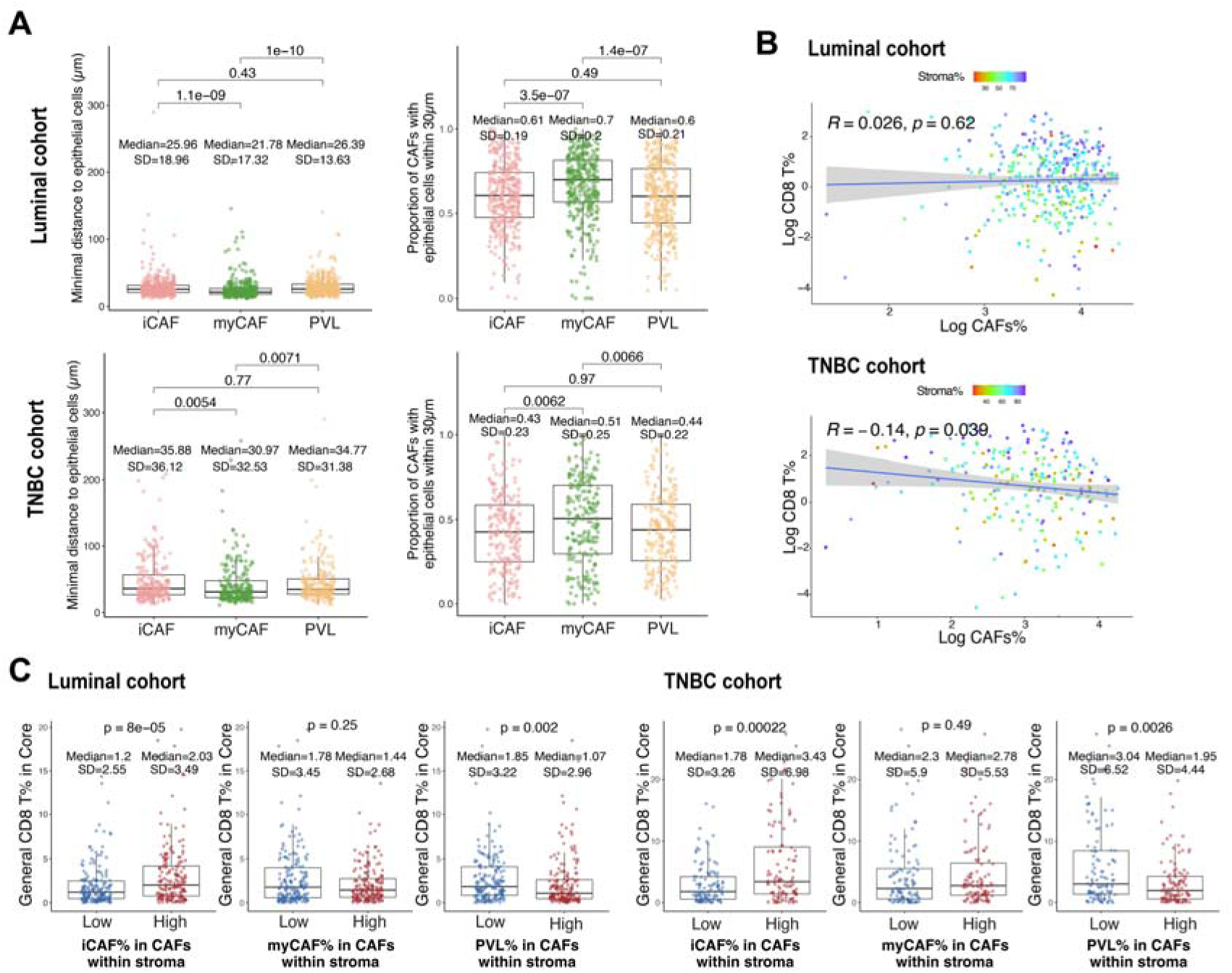
Spatial distribution of stromal cell types and their correlation with CD8 T cell infiltration. **A.** Median distance from iCAFs, myCAFs, PVLs to their nearest epithelial cells, and the proportion of each stromal cell type with epithelial cells present within a 30 µm radius. **B.** Correlation between the log-transformed proportions of CD8^+^ T cells and total stromal cells. **C.** PD1^-^CD8^+^ T cell percentages in TMA cores. Cores are stratified into low and high groups based on the median proportions of myCAFs, PVLs, and iCAFs within the total stromal population in the stromal region.

We observed no significant association between overall CAF abundance within the stromal compartment and T-cell infiltration in luminal cases (Pearson’s R = 0.026, p = 0.62) (**Figure 2B**). In TNBC, however, there was a modest negative correlation (Pearson’s R = –0.14, p = 0.039, **Figure 2B** When stratifying by stromal subset, divergent associations with CD8^+^ T cell infiltration emerged. Cores enriched in PVL cells exhibited reduced CD8^+^ T cell infiltration (p=0.002 for luminal; p=0.0026 for TNBC; **Figure 2C**). Conversely, iCAF-enriched cores demonstrated increased CD8^+^ T cell infiltration (p<0.001 for luminal; p<0.001 for TNBC, **Figure 2C**), indicating a possible role for iCAFs in supporting or permitting T cell entry into the TME.

### Immune compositions and spatial distribution reveal three immune infiltration phenotypes

To further characterise immune infiltration patterns in breast cancer, we classified TMA cores into three immune phenotypes based on T cell enrichment and tumour–T cell co-localisation (**Figures 3A, B**). The Immune cold phenotype was defined by total CD8^+^ T cell proportions below the median across all TMA cores (1.44% in luminal and 1.36% in TNBC). Immune segregated cores displayed spatial separation between tumour and T cell aggregates (cross-K AUC < 0, CD8^+^ T% ≥ 1.44% in luminal, CD8^+^ T% ≥ 1.36% in TNBC). Immune intermixing cores exhibited co-localisation of tumour cells and CD8^+^ T cells (cross-K AUC ≥ 0, CD8^+^ T% ≥ 1.44% in luminal, CD8^+^ T% ≥ 1.36% in TNBC).

**Figure 3.**
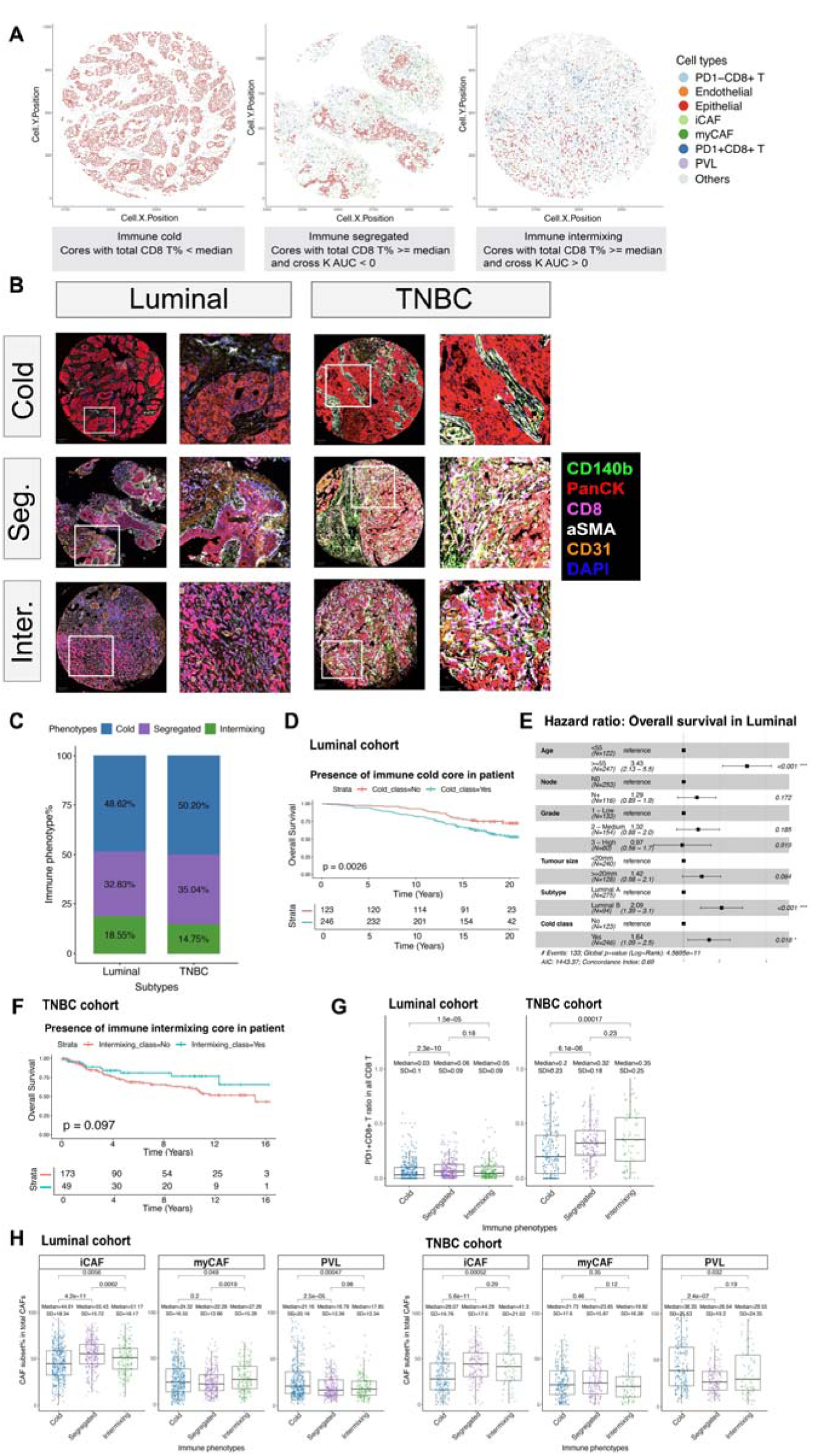
Immune phenotypes defined by T cell proportions and the co-localisation between T cells and epithelial cells. **A.** Cell type maps of represented cores from the luminal cohort with immune cold, immune segregated and immune intermixing phenotypes. **B.** mIF images of three immune phenotypes in luminal and TNBC cohorts. Image on the left shows the entire TMA core; image on the right shows a zoom-in region. The white box indicates the location of the zoom-in region in the TMA core. **C.** Percentages of immune phenotypes in luminal and TNBC cohorts. **D.** KM curves for OS stratified by the presence of immune cold cores in the luminal cohort. **E.** Multivariable Cox proportional hazard regression analysis of OS for the presence of immune cold cores in the luminal cohort, considering age, lymph node metastasis, grade, tumour size, and molecular subtypes. **F.** KM curves for OS stratified by the presence of immune intermixing cores in the TNBC cohort. **G.** The ratio of PD1^+^CD8^+^ T cells to all CD8^+^ T cells within immune cold, segregated, and intermixing cores. **H.** Percentages of iCAFs, myCAFs, and PVLs in total stromal cells within TMA cores of different immune phenotypes.

The most prevalent phenotype was immune cold in both cohorts, with a slightly higher proportion in TNBC (48.62% in luminal, 50.20% in TNBC). This was followed by immune segregated (32.83% in luminal, 35.04% in TNBC) and immune intermixing (18.55% in luminal, 14.75% in TNBC; **Figure 3C**). The presence of immune cold cores was associated with reduced survival in luminal tumours but not in TNBC (luminal: univariate p=0.0026; multivariate p=0.018, **Fig. 3D, E**). While in TNBC, the presence of intermixing cores showed a trend to be associated with better survival (p=0.097, **Figure 3F**). It is worth noting that an increased CD8^+^ T cell% was associated with better survival in the luminal cohort (univariate p=0.027), but not significant when accounting for clinical variables (multivariate p=0.211). This suggests that the presence of an immune cold region better predicts the patient outcome than a summarised proportion of PD1^+^CD8^+^ or PD1^-^CD8^+^ T cells. PD1^+^CD8^+^ T cell ratios display a similar level between segregated and intermixing cores in both subtypes, which were higher than that in the immune cold cores (**Fig. 3G**). The composition of CAF subsets varied by immune phenotype. In both cohorts, iCAFs were most abundant in immune segregated and intermixing phenotypes, and least abundant in immune cold cores (**Figure 3H**). In contrast, PVL cells were most abundant in immune cold cores (**Figure 3H**). myCAFs were more uniformly distributed, with slightly higher levels in intermixing phenotypes in luminal cases (**Figure 3H**). These findings suggest that distinct stromal subsets influence T-cell infiltration in different manners. PVLs may play a role in excluding T cells from infiltrating the tumour whereas iCAFs may relate to separation of cancer and T cells.

### Endothelial-distal PVLs are associated with T-cell exclusion

PVL is defined as a perivascular-like stromal cell population marked by CD146 positivity, these include pericytes, vascular smooth muscle cells and other mural cells. In previous studies, we observed that a subset of PVLs is spatially dissociated from endothelial cells, suggesting that these cells may have functions distinct from their conventional role in vascular contractility and blood flow regulation (9). In the luminal cases, we found that about 97.12% of PVLs were distant (>30 µm) from endothelial cells. Even at a 100⍰µm threshold, a large majority of PVLs remained distant from endothelial cells (**Figure 4A, B**). TNBC exhibited a lower proportion of disseminated PVLs (70.12% at 30 µm, 22.11% at 100 µm, **Supplementary figure 2A**), likely due to the higher endothelial abundance in TNBC compared to luminal subtype (**Figure 1G**). There was no prominent disseminated PVL in normal breast samples, suggesting the dissociation of PVLs from endothelial cells is specific to tumours (**Figure 4B**).

**Figure 4.**
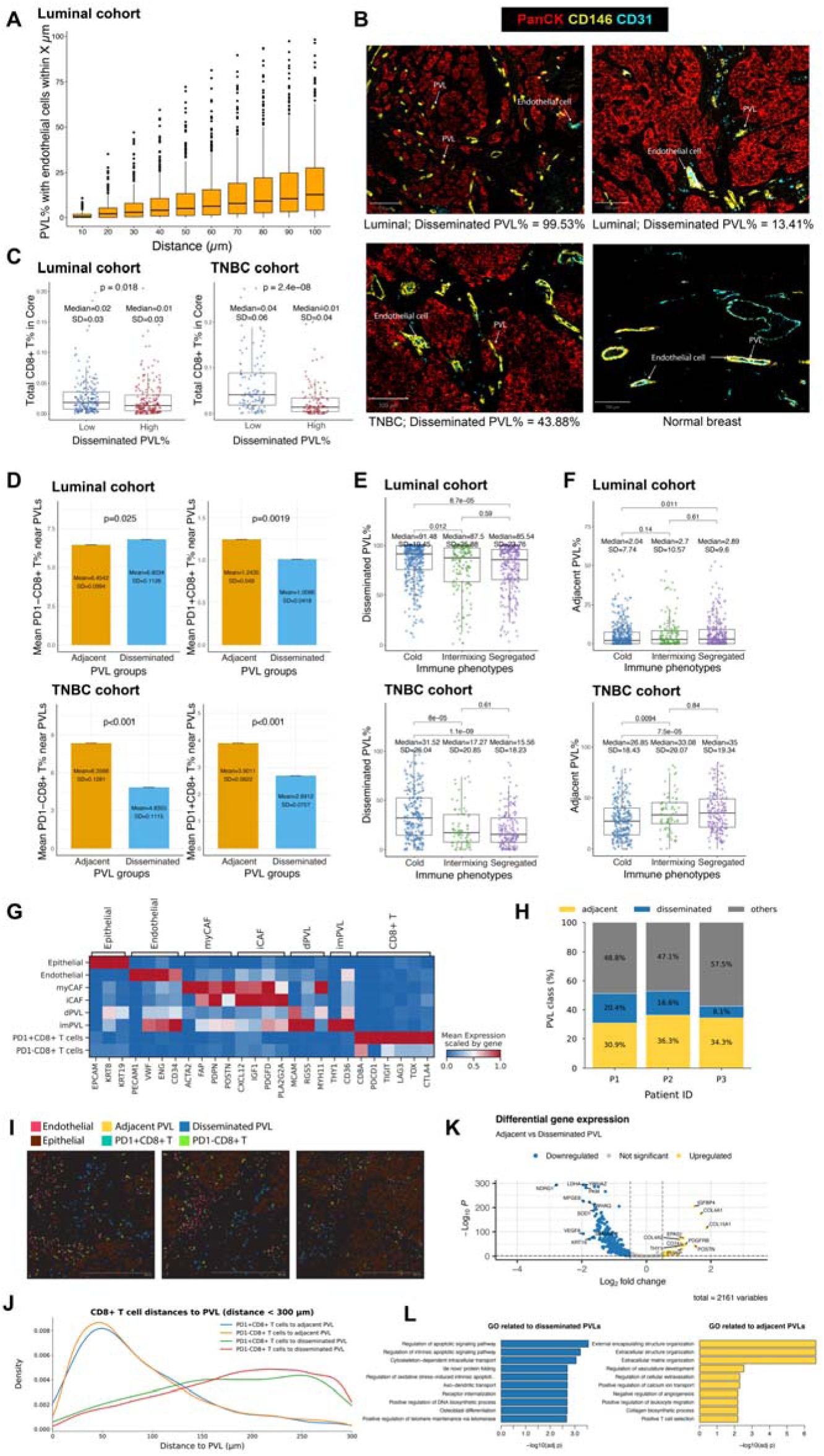
PVLs distal from endothelial cells are associated with T cell exclusion. PVLs were classified as adjacent if an endothelial cell was present within 30⍰µm, and as disseminated if no endothelial cell was present within 100⍰µm. **A.** Percentage of PVLs with endothelial cells present within a 10–100 µm radius in the luminal cohort. **B.** mIF images of TMA cores exhibiting disseminated PVLs in luminal and TNBC tumours, and the co-localisation of PVL and blood vessels in a normal breast sample. **C.** Percentages of CD8^+^ T cells in TMA cores with high and low disseminated PVL% stratified by the median. **D.** Mean percentages of PD1^+^CD8^+^ and PD1^-^CD8^+^ T cells in the neighbourhood of adjacent and disseminated PVLs. Neighbourhood is defined as the 10 nearest cells to each PVL. Error bar indicates the standard error. **E, F.** Percentages of disseminated and adjacent PVLs in the stromal region of cores classified as immune cold, segregated, or intermixing phenotypes. **G.** Gene expression heatmap of canonical marker genes across cell types identified in Xenium data. Cell types were defined using the same combination of markers derived from the mIF panel. Expression values represent mean counts scaled per gene. **H.** Bar plots showing the proportions of adjacent, disseminated, and other PVLs across Xenium sections from three TNBC tumours. **I.** Representative regions from Xenium sections illustrating the spatial distribution of PVLs, endothelial cells, epithelial cells, and CD8^+^ T cells. Both PD1^+^CD8^+^ and PD1^-^CD8^+^ T cells were found near adjacent PVLs, whereas far away from disseminated PVLs. **J.** Density plot of PD1^+^CD8^+^ and PD1^-^CD8^+^ T cells across a distance (0-300μm) to the nearest adjacent or disseminated PVL. **K.** Volcano plot of DEGs between adjacent and disseminated PVLs. DEGs with an absolute log_2_ fold change > 0.5 and adjusted P-value < 0.05 were classified as significantly up- or downregulated. The top 10 DEGs by log_2_ fold change in each direction were labelled. **L.** Bar plots showing the top 10 enriched GO Biological Process terms for the top 100 genes upregulated in disseminated PVLs (left, blue) and adjacent PVLs (right, yellow). Bar length represents the −log_10_ adjusted P-value, with greater values indicating stronger enrichment.

To investigate whether disseminated PVLs form at the expense of PVL-bounded vessels, we assessed the correlation between disseminated PVL (PVLs without endothelial cell present within 100 µm) and the proportion of endothelial cells with PVL present within 30 µm. No significant association was observed (luminal, Pearson’s R=-0.037, p=0.51; TNBC, Pearson’s R=0.077, p=0.26, **Supplementary figure 2B, C**). This suggests the existence of a spatially distinct PVL population that is not vessel-dependent.

We have observed a reduced CD8^+^ T cell abundance in PVL-enriched tumours (**Fig. 2C**). To explore the relationship between PVL spatial distribution and T cell infiltration, we examined the association between disseminated PVL abundance and CD8^+^ T cell proportions across cores. In luminal and especially in TNBC cohorts, CD8^+^ T cell infiltration was reduced in cores with a higher percentage of disseminated PVLs, suggesting that this PVL subpopulation may negatively regulate T cell entry (luminal: p=0.018; TNBC: p<0.001, **Figure 4C**). To better understand this, we compared the cellular neighbourhoods surrounding PVLs distal versus adjacent to endothelial cells. In both luminal and TNBC, disseminated PVLs had lower proportions of PD1^+^CD8^+^ T cells in their vicinity compared to adjacent PVLs (p<0.01, **Figure 4D**). This supports the observation that disseminated PVLs are associated with T cell exclusion. As further evidence, the proportion of disseminated PVLs in the stromal region was highest in immune cold cores across both subtypes (**Figure 4E**). In contrast, adjacent PVLs were more abundant in segregated and intermixing subtypes (**Figure 4F**). Together, these findings suggest that the influence of PVLs on T cell exclusion was primarily associated with vessel independent PVLs.

To validate the immune-cold association of disseminated PVLs, we generated an independent spatial transcriptomics dataset for three TNBC tumours using the Xenium Prime platform (5001 plus 100 genes add-on panel, **Supplementary table 4**). Stromal and immune phenotypes were annotated based on the combined expression of genes corresponding to the markers used in the mIF panel (**Supplementary table 5**). The classified cell types exhibited expression of canonical lineage markers, confirming the fidelity of our cell type annotation (**Figure 4G**) (22). The overlap of *VWF, ENG*, and *CD34* for endothelial cells and imPVL can be attributed to the spill-over of transcripts from endothelial cells. Within this dataset, disseminated PVLs accounted for an average of 15% of the total PVLs (range: 8.1% to 20.4%), comparable to the prevalence observed in the mIF TNBC cohort (**Figure 4H**). Spatial analysis revealed that both PD1^-^CD8^+^ and PD1^+^CD8^+^ T cells were distributed further away from disseminated PVLs than from adjacent PVLs (**Figure 4I, J**), consistent with the CD8^+^ T cell exclusion associated with disseminated PVLs in the mIF TNBC cohort. Differential gene expression analysis revealed upregulation of genes related to extracellular matrix organisation and vascular stability in adjacent PVLs, including *COL15A1, COL4A1, POSTN, IGFBP4*, and *PDGFRB* (**Figure 4K, L**). In contrast, disseminated PVLs exhibited upregulation of genes related to vascular permeability, hypoxia signalling and apoptosis, such as *NDRG1, VEGFA, LDHA, MFGE8*, and *YWHAZ* (**Figure 4K, L**) (23–25). These divergent expression profiles suggest that disseminated PVLs transition away from a vessel maintenance role towards a stress-response state, potentially linked to aberrant vasculature and reduced immune cell infiltration.

### iCAFs constrain spatial contacts between T cells and cancer cells in luminal breast cancer

In luminal tumours, iCAFs were enriched in immune segregated and intermixing cores, indicating that their presence is not directly associated with an immune cold phenotype (**Figure 3H, 5A**). However, we observed that the minimal distance between CD8^+^ T cells and epithelial cells in luminal cancers was greater in the presence of iCAFs (**Figure 5B**), suggesting that iCAFs may spatially restrict T cells within the stromal compartment.

**Figure 5.**
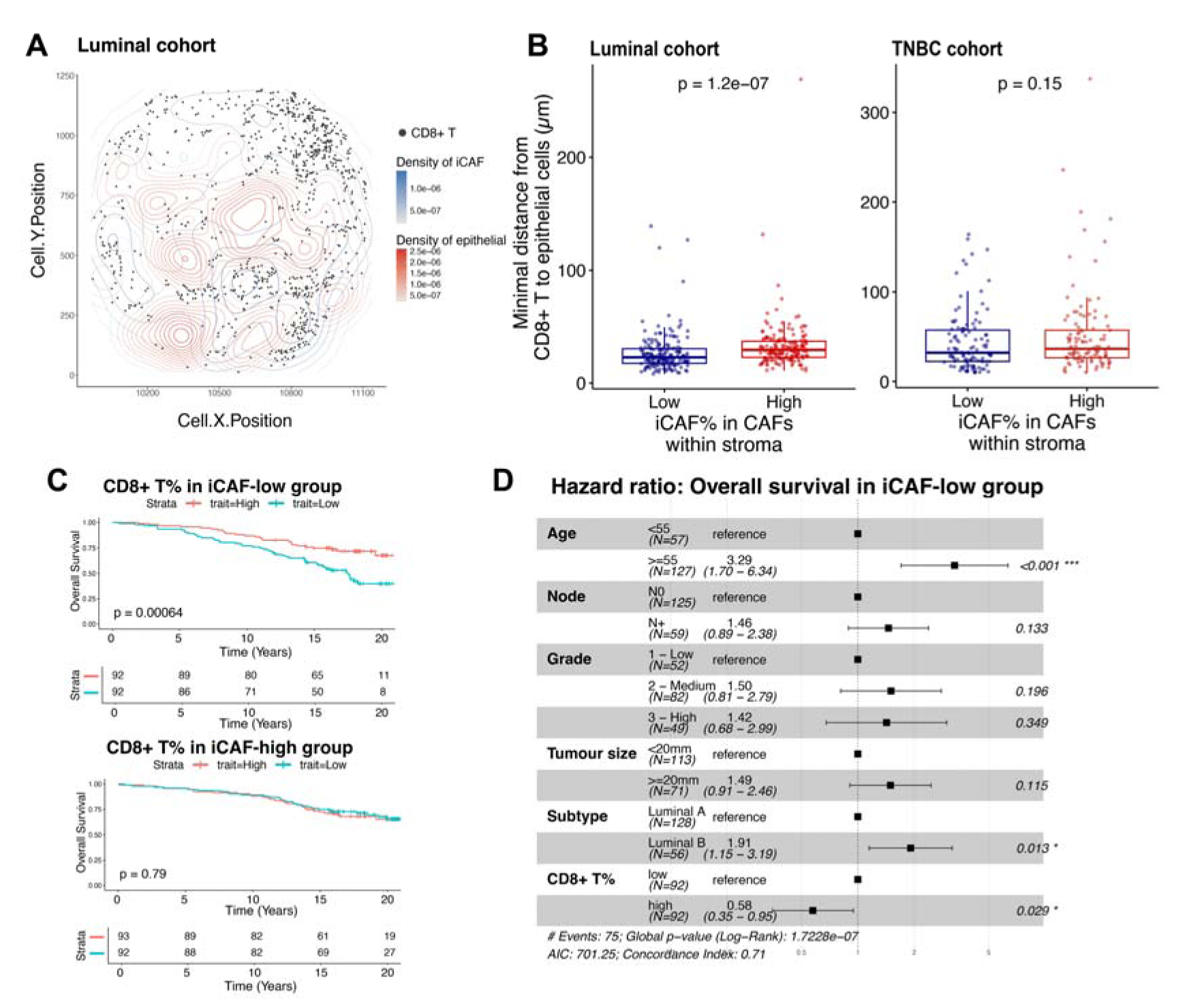
iCAF are associated with the segregation of CD8 T cells from tumour in the luminal cohort. **A.** Representative TMA core showing the spatial distribution of CD8⁺ T cells in iCAF-dense (blue) and epithelial-dense (red) regions. **B.** Minimal distance from CD8⁺ T cells to the nearest epithelial cell in TMA cores with low vs. high iCAF proportions. **C.** KM curves for OS stratified by the percentage of PD1⁻CD8⁺ T cells in iCAF-high vs. iCAF-low groups. **D.** Multivariable Cox proportional hazard regression analysis of OS for the proportion of CD8⁺ T cells% in iCAF-low group, considering age, lymph node metastasis, tumour grade, tumour size, and molecular subtype.

Remarkably, CD8^+^ T cell infiltration was associated with improved patient survival for luminal tumours with low iCAF abundance in both univariate and multivariate analysis (**Figure 5C, D**), but this prognostic value was lost in iCAF-high tumours. This suggests that high iCAF was associated with T cell dysfunction. Similarly, CD8^+^ T cell infiltration was associated with improved survival in myCAF-low tumours but not in myCAF-high tumours (**Supplementary figure 3A**). However, the significance was lost after accounting for clinical variables, especially age at diagnosis (**Supplementary figure 3B**). One possible explanation is that myCAFs may also be associated with T cell dysfunction, although we were unable to explore this further due to limitations of our dataset.

### Clinical correlation of spatially resolved cell compositions

To assess the correlation of spatial context of stromal subsets with clinical characteristics, we evaluated the enrichment of six spatial features per stromal subset in patients stratified by prognostic variables from survival analysis: age at diagnosis, nodal metastasis, tumour grade, histological subtypes, molecular subtypes and TILs (**Supplementary table 6, 7**). The spatial features include the proportion of PVL, myCAF, and iCAF within the stromal compartment, together with their spatial proximity to endothelial, epithelial, and CD8 T cells measured by minimal distance, normalised mixing score, and proportion of adjacent cells (Methods, **Supplementary table 3**). Strong association with clinical variables were defined by significant elevation of at least two spatial features within the same clinical group.

In luminal tumours, increased proximity between PVL and cancer epithelial cells, as reflected by a higher proportion of cancer-adjacent PVLs and shorter PVL-epithelial minimal distance, was linked to lymph node metastasis (**Figure. 6A**). Enhanced iCAF-cancer proximity showed similar associations with lymph node metastasis (**Figure. 6A**), whereas myCAF-cancer adjacency did not show significant relationship with assessed clinical variables (**Supplementary figure 4A**) On the other hand, closer proximity between iCAF and CD8^+^ T cells was observed for tumours diagnosed at a younger age (Age<55, **Figure 6A**). There was a similar trend for PVL-CD8 T cell proximity but were significant for the minimal distance only. Additionally, increased stromal iCAF proportions distinguished luminal A subtype and low-grade tumours (**Figure 6A**). For all three stromal cell types, their adjacency to endothelial cells increased in high-grade tumours (**Figure 6A**, **Supplementary figure 3A**). This is due to the enrichment of endothelial cells in high-grade luminal cases (**Supplementary figure 4B**).

**Figure 6.**
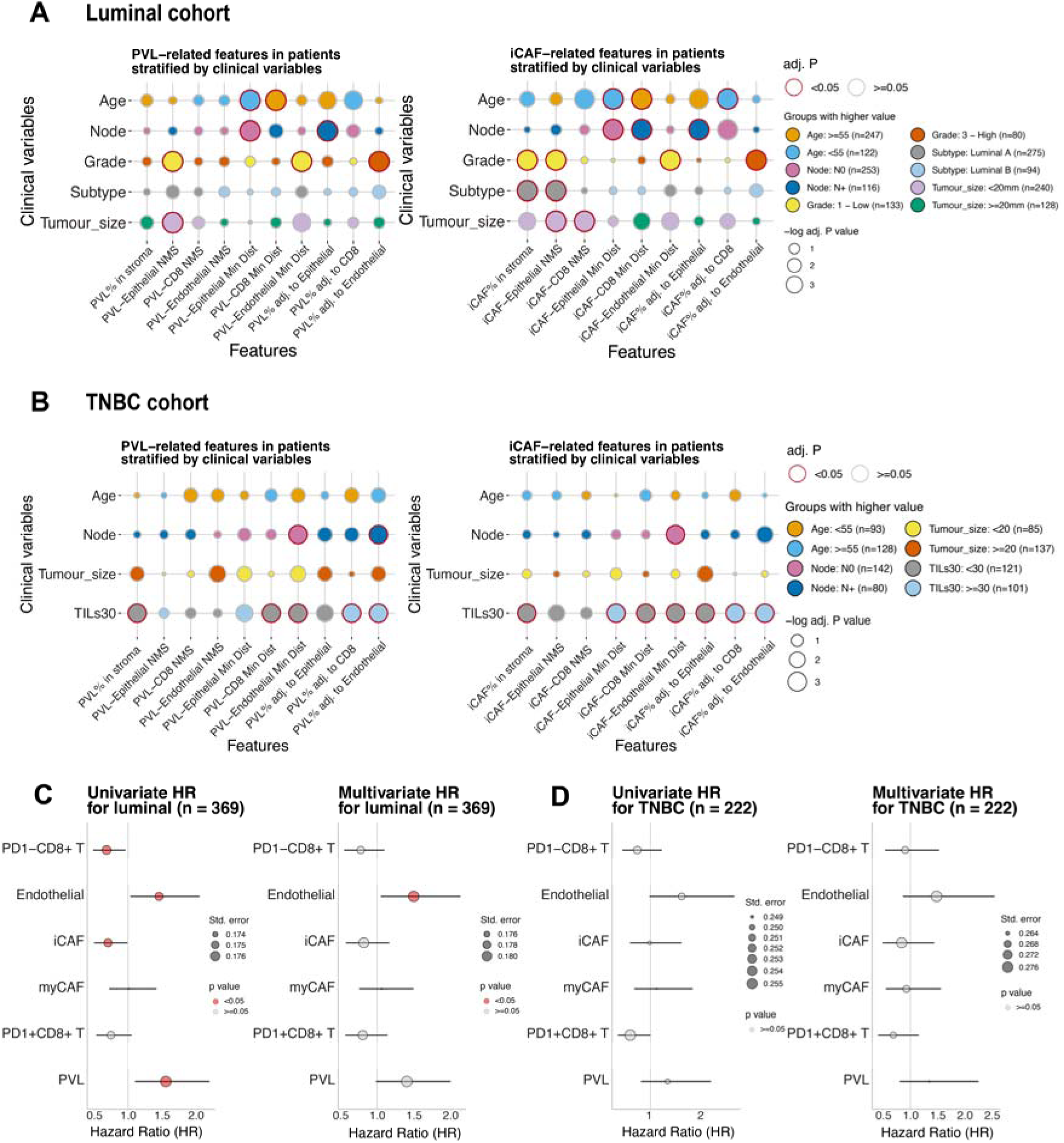
Association of iCAF- and PVL-related spatial features with clinical variables in the luminal and TNBC cohorts. Patients were stratified by clinical variables. Dot colours indicate patient groups in which the feature is increased. Dot outlines denote statistical significance of feature enrichment (red: significant; grey: not significant), assessed by the Wilcoxon rank-sum test with p-values corrected using the BH method. NMS, normalised mixing score. **C.** Univariate and multivariate analysis of the association between cell type proportions and OS in the luminal cohort. **D.** Same as **C** for the TNBC cohort.

In TNBC, PVL-endothelial proximity correlated with nodal metastasis (**Figure. 6B**). Across all the three stromal cell types, high proportions in the stroma and increased spatial separation from CD8^+^ T cells and endothelial cells associated with reduced immune infiltration (**Figure 6B**, **Supplementary figure 4C**). Additionally, enhanced iCAF and myCAF adjacency to cancer cells corresponded with reduced TILs (**Figure 6B**, **Supplementary figure 4C**). These findings indicate that in TNBC, CD8^+^ T cell and endothelial cell enrichment strongly correlate with clinical TIL assessment, whereas stromal cell abundance generally inversely correlates with immune infiltration.

In terms of patient outcomes, we stratified cohorts into ‘high’ or ‘low’ groups based on median cell-type proportions across patients. In the luminal cohort, reduced PD1^-^CD8^+^ T cells and iCAFs proportions, and high PVLs and endothelial cell proportions were associated with inferior OS (**Figure 6C**). In multivariate analysis adjusting for age, node metastasis, tumour grade, tumour size and molecular subtype, endothelial cell abundance remained significantly associated with worse survival (**Figure 6C**). In TNBC, similar trends were demonstrated for PD1^-^CD8^+^ T cells, PVLs and endothelial cells, but none of the cell types achieved statistical significance in univariate or multivariate analysis (**Figure 6D**).

## DISCUSSION

In this study, we have characterised the spatially distinct features of stromal subsets in relation to immune cells (**Figure 7**). This was done in two large breast cancer cohorts consisting of over 1,300 TMA cores from 591 patients. Both cohorts have well-annotated clinicopathological data as well as robust long-term outcome data with follow-up of up to 16 years. This gives our study sufficient power to identify clinically meaningful correlations, while addressing limitations of many prior spatial profiling studies that relied on only a small number of samples or patients and were therefore constrained by tissue selection biases and the challenges of both intra-tumoural and inter-tumoural heterogeneity.

**Figure 7.**
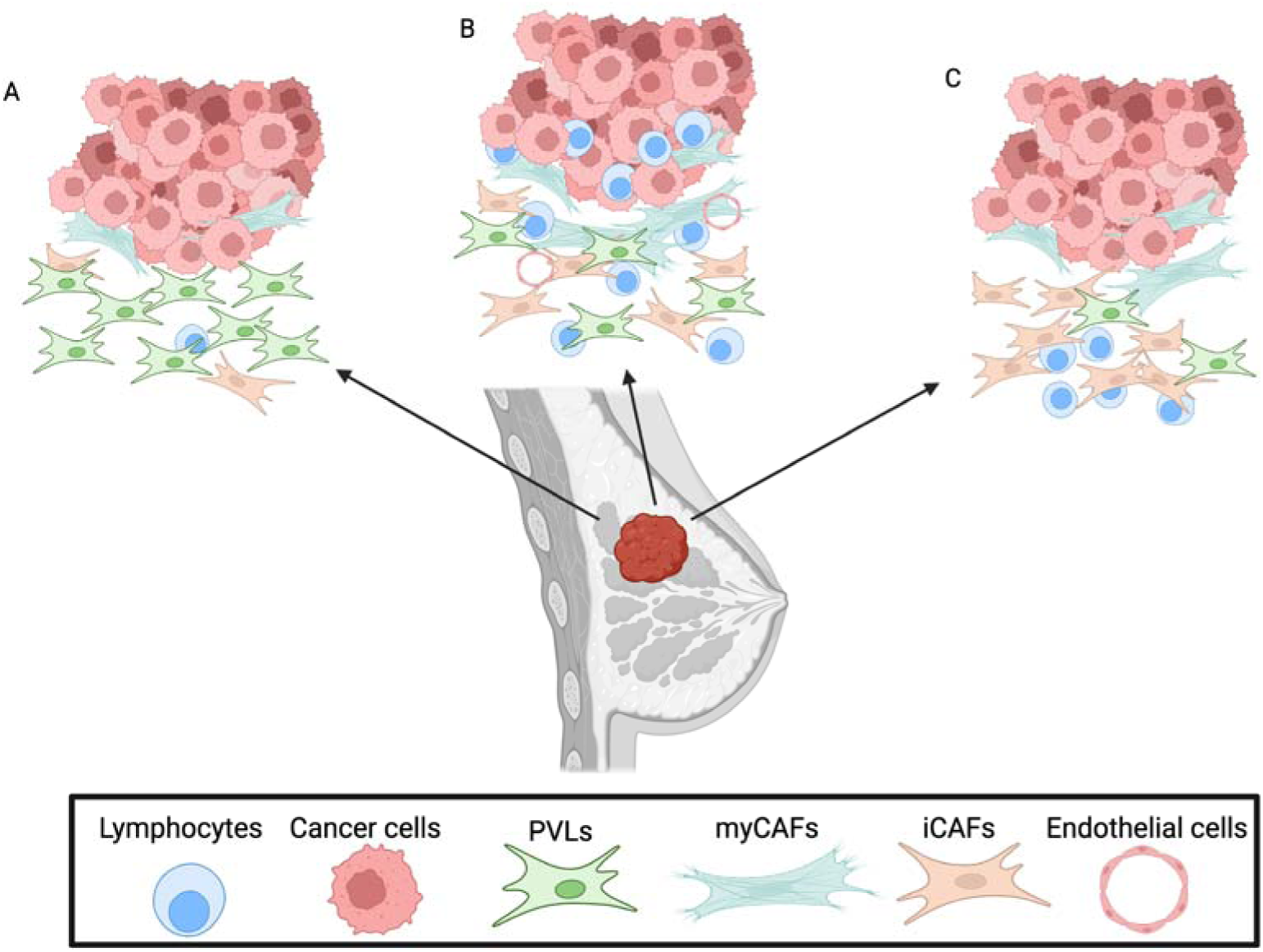
Diagrammatic summary of key findings. (Created in BioRender. Chen, J. (2026) https://BioRender.com/d28tsrm). **A.** Disseminated PVLs are associated with T cell exclusion and are enriched in immune cold phenotype. **B.** myCAFs are located in close proximity to cancer cells, iCAFs and PVLs are disseminated throughout the stroma. **C.** iCAFs are enriched in immune segregated phenotype where T cells are present but restricted away from cancer cells.

We make the novel observation that the large majority of PVLs are located distant from endothelial cells, despite their perivascular lineage (**Figure 7A**). In breast cancers, but not normal breast, the majority of PVLs were disseminated throughout the stroma rather than restricted to vessel-adjacent regions. Morphologically these cells closely resemble CAFs and share expression of canonical CAF markers such as ACTA2 and PDGFRβ and most likely overlap with cells annotated as vascular-CAFs (vCAFs) in prior studies (3, 22). In Bartoschek et al. (3), vCAFs were defined as cells enriched for mural genes such as *NOTCH3, EPAS1, COL18A1 and NR2F2* and perivascular markers such as *RGS5, MCAM, DES, KCNJ8 and HIGD1B*. They do not uniformly exhibit classical fibroblast markers such as *DCN, FAP* and *PDGFRα*. It was also observed that these cells lay along vessels early and become increasingly detached with tumour progression (3), consistent with perivascular origin and delamination. At the protein level, vCAFs were fined as CD146^high^ CD31^neg^ with FAP and PDPN used as discriminators for ‘activated CAFs’ and pericytes required extra markers such as RGS5 (22). vCAF and pericytes were hard to distinguish in some datasets in this study, underscoring phenotypic overlap between these two subsets. Our previously published scRNA-seq data demonstrated distinct transcriptional differences between PVLs and CAFs, supporting their unique identity within the stromal compartment. Gene expressions from our Xenium data demonstrated the consistency between the CAF and PVL cell types identified by mIF and their corresponding transcriptomic profiles (**Figure 4G**). Together, our studies and the literature support recasting ‘vCAFs’ as perivascular-like stromal cells that frequently lack classical fibroblast-ECM features and may delaminate from vessels into the tumour stroma. In normal breast tissue, this PVL subset resembles vascular smooth muscle cells, transcriptionally and spatially distinct from pericytes (26).

Our study is the first to link increased PVL proportion with CD8^+^ T cell exclusion and a cold immunophenotype (**Figure 7A**), consistent across both luminal and TNBC cohorts. This discovery was uniquely enabled by the application of a marker panel targeting stromal and lymphocyte subsets to large cohorts of disease. In the luminal cohort, higher PVL proportion was prognostic of poorer OS. While this was not statistically significant in the TNBC cohort, potentially limited by its smaller sample size, a similar trend was observed. PVL-associated T cell exclusion was primarily associated with the subpopulation of disseminated PVLs, where increased endothelial-distal PVLs correlated with decreased CD8^+^ T cell infiltration. Spatial transcriptomic analysis revealed that both PD1^+^CD8^+^ and PD1^-^CD8^+^ T cells were localised further away from disseminated PVLs while closer to adjacent PVLs, confirming the association of disseminated PVLs with immune cold phenotype and T cells exclusion (**Figure 4J**). Additionally, disseminated PVLs displayed an upregulation of genes associated with vascular permeability and hypoxia response, which is a new insight into these cells (**Figure 4L**). Previous studies examining endothelial-pericyte interactions have observed that pericyte detach and migrate away from blood vessels as a response to aberrant PDGFβ signalling (27, 28). These detached pericytes can then differentiate into myofibroblasts, contributing to fibrosis (29, 30), and also lead to vessel instability and vulnerability (31). Our disseminated PVLs may represent these detached pericytes, however, our data suggests these cells do not differentiate into myCAF and maintain CD146 expression. More work is warranted to investigate signalling pathways driving these phenotypes, which may potentially be targeted to reverse associated effect on T cell infiltration.

myCAFs and iCAFs have been extensively reported in the literature. Consistent with previous studies (8, 32, 33), we found that myCAFs were located in close proximity to epithelial cells whereas iCAFs were more broadly distributed throughout the stroma (**Figure 7B**). Our study further suggests that iCAFs may spatially restrict T cells within the stromal compartment, as evidenced by an association of iCAFs with a segregated immunophenotype (**Fig 3H**) and increased distance between CD8^+^ T cell and epithelial cells in the presence of iCAFs (**Figure 7C**). This is further supported by the diminished survival benefit conferred by T cell infiltration in iCAF-high tumours (**Figure 5C**). We propose that this may be mediated by chemokine directed migration of T cells towards iCAFs, which robustly express the T cell chemokines CXCL12 and CCL2 (8).

Spatial immunophenotypes have been shown to correlate with survival and response to immune checkpoint inhibitors in breast cancer (34–36). By classifying our cores into immune cold, segregated and intermixing phenotypes, we demonstrated a significantly poorer survival in patients with the presence of immune cold cores in luminal tumours, adding to existing evidence that spatial immune contexture is prognostic in breast cancer. Furthermore, we examined stromal phenotypes in these immunophenotypes and found an enrichment of PVLs in immune cold cores, consistent with our finding that PVLs correlate with T cell exclusion (**Figure 7A**). This was primarily related to endothelial-distal PVLs whereas endothelial-adjacent PVLs were not enriched in immune cold cores. This also suggests that this effect was not attributable to blood vessel density. We found that iCAFs were enriched in immune segregated cores where T cells were present but restricted to the tumour periphery (**Figure 7C**). T-cell exclusion is one of the leading causes of resistance to immune checkpoint inhibition (37) and recent studies suggest a role of CAFs in mediating this (38, 39). Our result provides further evidence to support this and suggests that PVLs and iCAFs contribute to this in different ways. While PVLs exclude T cells from migrating into the tumour, iCAFs were related to the segregation of cancer and T cells. While our findings provide novel observations on how stromal heterogeneity, particularly PVLs and iCAFs, shape CD8^+^ T cell infiltration within a spatially relevant context and demonstrates its clinical relevance, further work is required to elucidate the functional and mechanistic basis underlying these observations.

For decades, conventional methods such as immunohistochemistry have been the primary tools for tumour characterisation. However, these methods are limited in their capacity to capture the complexity of cellular diversity within tumours. The application of mIF and digital pathology enables the use of expanded protein panels, providing a more comprehensive, spatially resolved view of tumours and their TME. However, these technologies require greater standardisation and automation of staining and imaging protocols to enhance reproducibility and clinical applicability (12). One of the limitations of our findings is the potential overcalling of iCAFs due to the lack of a positive marker for iCAF in our mIF panel. iCAFs were annotated as PDGFRβ^+^ cells negative for other stromal markers (αSMA, CD146, THY1) and hence may include non-iCAF phenotypes belonging to subsets not annotated by these markers, such as antigen-presenting CAFs and matrix CAFs (2).

This study is based on TMA cores rather than whole slide sections to improve the feasibility to analyse large clinical cohorts to strengthen potential clinical applicability and statistical power. To account for the well-documented intra-tumoural heterogeneity of breast cancer, we utilised a sampling strategy of three representative 1mm cores. This approach is supported by validation studies demonstrating that three-core sampling provides high concordance with whole-tissue sections for standard biomarkers and clinical outcomes, capturing the diversity of the TME while maximising experimental throughout for spatial transcriptomic analysis (40, 41). Results of our Xenium profiling of surgical resections demonstrated similar findings in the prevalence of disseminated PVLs and their association with CD8^+^ T cell exclusion, providing further reassurance that our findings are not substantially affected by the use of TMA cores. While the QuPath algorithm worked well to allow objective quantitative cell annotation suitable for application to large clinical cohorts with efficient high throughput of data, the training of slides was semi-quantitative, subject to interobserver variability. We addressed this by manually cross-checking cores across different slides, supervised by a pathologist. Advancement in deep learning artificial intelligence algorithm will help to overcome these technical challenges to allow automated deep interrogation of vast quantities of data generated from whole tumours of large cohorts.

## CONCLUSION

Using mIF to map the stromal-immune dynamics in breast cancer in two large clinical cohorts, our study highlights that stromal subsets exhibit distinct spatial distribution, interaction with immune cells and clinical correlations. The clinical correlations identified in our study warrant further investigation as potential biomarkers, ideally through incorporation into prospective clinical trials. Specific stromal-immune interactions may also be explored as novel therapeutic targets to modulate the tumour immune microenvironment and enhance responses to immunotherapy.

## Acknowledgements

J.C. received scholarships from the UNSW Scientia program and SPHERE Cancer Clinical Academic Group. A.S. is a Breast Cancer Research Foundation investigator, the Peter Chair in Breast Cancer Research, and is supported by the generosity of John McMurtrie, AM and Deborah McMurtrie. E.L. is an endowed chair of National Breast Cancer Foundation Australia. This work was supported by a grant from the National Breast Cancer Foundation Australia, grant RG22⍰10 from Cancer Council NSW, and grants from NHMRC.

## Author contributions

J.C.: project design, methodology, image and data analysis, manuscript writing and review H.Z.: data analysis, manuscript writing and review T.R.: supervision, manuscript review S.W.: project design, methodology I.S.: methodology, image acquisition and analysis E.M.: sample acquisition, methodology, supervision, manuscript review P.G./J.L./L.B.: sample acquisition, clinical data collection, manuscript review E.L.: project design, resources, supervision, funding provision, manuscript review A.S.: project design, resources, supervision, manuscript review.

## REFERENCE

1. Bray F, Laversanne M, Sung H, Ferlay J, Siegel RL, Soerjomataram I, et al. Global cancer statistics 2022: GLOBOCAN estimates of incidence and mortality worldwide for 36 cancers in 185 countries. CA Cancer J Clin. 2024;74(3):229–63.

2. Chhabra Y, Weeraratna AT. Fibroblasts in cancer: Unity in heterogeneity. Cell. 2023;186(8):1580–609.

3. Bartoschek M, Oskolkov N, Bocci M, Lovrot J, Larsson C, Sommarin M, et al. Spatially and functionally distinct subclasses of breast cancer-associated fibroblasts revealed by single cell RNA sequencing. Nat Commun. 2018;9(1):5150.

4. Elyada E, Bolisetty M, Laise P, Flynn WF, Courtois ET, Burkhart RA, et al. Cross-Species Single-Cell Analysis of Pancreatic Ductal Adenocarcinoma Reveals Antigen-Presenting Cancer-Associated Fibroblasts. Cancer Discov. 2019;9(8):1102–23.

5. Lambrechts D, Wauters E, Boeckx B, Aibar S, Nittner D, Burton O, et al. Phenotype molding of stromal cells in the lung tumor microenvironment. Nat Med. 2018;24(8):1277–89.

6. Puram SV, Tirosh I, Parikh AS, Patel AP, Yizhak K, Gillespie S, et al. Single-Cell Transcriptomic Analysis of Primary and Metastatic Tumor Ecosystems in Head and Neck Cancer. Cell. 2017;171(7):1611–24 e24.

7. Sebastian A, Hum NR, Martin KA, Gilmore SF, Peran I, Byers SW, et al. Single-Cell Transcriptomic Analysis of Tumor-Derived Fibroblasts and Normal Tissue-Resident Fibroblasts Reveals Fibroblast Heterogeneity in Breast Cancer. Cancers (Basel). 2020;12(5).

8. Wu SZ, Al-Eryani G, Roden DL, Junankar S, Harvey K, Andersson A, et al. A single-cell and spatially resolved atlas of human breast cancers. Nat Genet. 2021;53(9):1334–47.

9. Wu SZ, Roden DL, Wang C, Holliday H, Harvey K, Cazet AS, et al. Stromal cell diversity associated with immune evasion in human triple-negative breast cancer. EMBO J. 2020;39(19):e104063.

10. Savas P, Virassamy B, Ye C, Salim A, Mintoff CP, Caramia F, et al. Single-cell profiling of breast cancer T cells reveals a tissue-resident memory subset associated with improved prognosis. Nat Med. 2018;24(7):986–93.

11. Elhanani O, Ben-Uri R, Keren L. Spatial profiling technologies illuminate the tumor microenvironment. Cancer Cell. 2023;41(3):404–20.

12. Chen J, Larsson L, Swarbrick A, Lundeberg J. Spatial landscapes of cancers: insights and opportunities. Nat Rev Clin Oncol. 2024;21(9):660–74.

13. Millar EK, Graham PH, O’Toole SA, McNeil CM, Browne L, Morey AL, et al. Prediction of local recurrence, distant metastases, and death after breast-conserving therapy in early-stage invasive breast cancer using a five-biomarker panel. J Clin Oncol. 2009;27(28):4701–8.

14. Wang J, Browne L, Slapetova I, Shang F, Lee K, Lynch J, et al. Multiplexed immunofluorescence identifies high stromal CD68(+)PD-L1(+) macrophages as a predictor of improved survival in triple negative breast cancer. Sci Rep. 2021;11(1):21608.

15. Feng Y, Yang T, Zhu J, Li M, Doyle M, Ozcoban V, et al. Spatial analysis with SPIAT and spaSim to characterize and simulate tissue microenvironments. Nat Commun. 2023;14(1):2697.

16. Virshup Iea. anndata: Access and store annotated data matrices. Journal of Open Source Software. 2024;9(101), 4371, 10.21105/joss.04371.

17. Palla G, Spitzer H, Klein M, Fischer D, Schaar AC, Kuemmerle LB, et al. Squidpy: a scalable framework for spatial omics analysis. Nat Methods. 2022;19(2):171–8.

18. Love MI, Huber W, Anders S. Moderated estimation of fold change and dispersion for RNA-seq data with DESeq2. Genome Biol. 2014;15(12):550.

19. Xu S, Hu E, Cai Y, Xie Z, Luo X, Zhan L, et al. Using clusterProfiler to characterize multiomics data. Nat Protoc. 2024;19(11):3292–320.

20. Cortes J, Rugo HS, Cescon DW, Im SA, Yusof MM, Gallardo C, et al. Pembrolizumab plus Chemotherapy in Advanced Triple-Negative Breast Cancer. N Engl J Med. 2022;387(3):217–26.

21. Schmid P, Cortes J, Dent R, Pusztai L, McArthur H, Kummel S, et al. Event-free Survival with Pembrolizumab in Early Triple-Negative Breast Cancer. N Engl J Med. 2022;386(6):556–67.

22. Cords L, Tietscher S, Anzeneder T, Langwieder C, Rees M, de Souza N, et al. Cancer-associated fibroblast classification in single-cell and spatial proteomics data. Nat Commun. 2023;14(1):4294.

23. Brown LF, Berse B, Jackman RW, Tognazzi K, Guidi AJ, Dvorak HF, et al. Expression of vascular permeability factor (vascular endothelial growth factor) and its receptors in breast cancer. Hum Pathol. 1995;26(1):86–91.

24. Joshi V, Stacey A, Feng Y, Kalita-de Croft P, Duijf PH, Simpson PT, et al. NDRG1 is a prognostic biomarker in breast cancer and breast cancer brain metastasis. J Pathol Clin Res. 2024;10(2):e12364.

25. Sharma P, Chida K, Wu R, Tung K, Hakamada K, Ishikawa T, et al. VEGFA Gene Expression in Breast Cancer Is Associated With Worse Prognosis, but Better Response to Chemotherapy and Immunotherapy. World J Oncol. 2025;16(1):120–30.

26. Kumar T, Nee K, Wei R, He S, Nguyen QH, Bai S, et al. A spatially resolved single-cell genomic atlas of the adult human breast. Nature. 2023;620(7972):181–91.

27. Schrimpf C, Teebken OE, Wilhelmi M, Duffield JS. The role of pericyte detachment in vascular rarefaction. J Vasc Res. 2014;51(4):247–58.

28. Stratman AN, Schwindt AE, Malotte KM, Davis GE. Endothelial-derived PDGF-BB and HB-EGF coordinately regulate pericyte recruitment during vasculogenic tube assembly and stabilization. Blood. 2010;116(22):4720–30.

29. Humphreys BD, Lin SL, Kobayashi A, Hudson TE, Nowlin BT, Bonventre JV, et al. Fate tracing reveals the pericyte and not epithelial origin of myofibroblasts in kidney fibrosis. Am J Pathol. 2010;176(1):85–97.

30. Lin SL, Kisseleva T, Brenner DA, Duffield JS. Pericytes and perivascular fibroblasts are the primary source of collagen-producing cells in obstructive fibrosis of the kidney. Am J Pathol. 2008;173(6):1617–27.

31. Caruso RA, Fedele F, Finocchiaro G, Pizzi G, Nunnari M, Gitto G, et al. Ultrastructural descriptions of pericyte/endothelium peg-socket interdigitations in the microvasculature of human gastric carcinomas. Anticancer Res. 2009;29(1):449–53.

32. Andersson A, Larsson L, Stenbeck L, Salmen F, Ehinger A, Wu SZ, et al. Spatial deconvolution of HER2-positive breast cancer delineates tumor-associated cell type interactions. Nat Commun. 2021;12(1):6012.

33. Ohlund D, Handly-Santana A, Biffi G, Elyada E, Almeida AS, Ponz-Sarvise M, et al. Distinct populations of inflammatory fibroblasts and myofibroblasts in pancreatic cancer. J Exp Med. 2017;214(3):579–96.

34. Hammerl D, Martens JWM, Timmermans M, Smid M, Trapman-Jansen AM, Foekens R, et al. Spatial immunophenotypes predict response to anti-PD1 treatment and capture distinct paths of T cell evasion in triple negative breast cancer. Nat Commun. 2021;12(1):5668.

35. Shiao SL, Gouin KH, 3rd, Ing N, Ho A, Basho R, Shah A, et al. Single-cell and spatial profiling identify three response trajectories to pembrolizumab and radiation therapy in triple negative breast cancer. Cancer Cell. 2024;42(1):70–84 e8.

36. Wang XQ, Danenberg E, Huang CS, Egle D, Callari M, Bermejo B, et al. Spatial predictors of immunotherapy response in triple-negative breast cancer. Nature. 2023;621(7980):868–76.

37. Sharma P, Hu-Lieskovan S, Wargo JA, Ribas A. Primary, Adaptive, and Acquired Resistance to Cancer Immunotherapy. Cell. 2017;168(4):707–23.

38. Feig C, Jones JO, Kraman M, Wells RJ, Deonarine A, Chan DS, et al. Targeting CXCL12 from FAP-expressing carcinoma-associated fibroblasts synergizes with anti-PD-L1 immunotherapy in pancreatic cancer. Proc Natl Acad Sci U S A. 2013;110(50):20212–7.

39. Mariathasan S, Turley SJ, Nickles D, Castiglioni A, Yuen K, Wang Y, et al. TGFbeta attenuates tumour response to PD-L1 blockade by contributing to exclusion of T cells. Nature. 2018;554(7693):544–8.

40. Camp RL, Charette LA, Rimm DL. Validation of tissue microarray technology in breast carcinoma. Lab Invest. 2000;80(12):1943–9.

41. Torhorst J, Bucher C, Kononen J, Haas P, Zuber M, Kochli OR, et al. Tissue microarrays for rapid linking of molecular changes to clinical endpoints. Am J Pathol. 2001;159(6):2249–56.

