## Supplementary tables and figures for "Stromal cell subsets modulate T-cell infiltration in early breast cancer"

**Supplementary table 1.** Antibodies used in mIF panel.

| **Antibody** | **Manufacturer and catalogue number** | **Epitope** | **Dilution** |
| --- | --- | --- | --- |
| panCK | Abcam ab27988 | OPAL650 | 1: 2000 |
| PDGFRβ (CD140b) | Abcam ab32570 | OPAL 540 | 1: 1000 |
| aSMA | Abcam ab5694 | OPAL780 | 1: 500 |
| CD146 | Abcam ab75769 | OPAL 570 | 1: 1250 |
| THY1(CD90) | Abcam ab133350 | OPAL 690 | 1: 4000 |
| CD8 | Invitrogen MA5-13473 | OPAL 620 | 1: 1000 |
| PD-1 | Abcam ab137132 | OPAL 520 | 1: 50 |
| CD31 | Agilent Technologies/DAKO M0823 | OPAL 480 | 1: 100 |

**Supplementary table 2.** Marker combinations for defining cell types in the mIF panel. Alternative combinations were separated by semi comma.

| **Cell type** | **Phenotype marker combinations** |
| --- | --- |
| myCAF | a-SMA+CD140b+ |
| Epithelial cells | PanCK+; CD31+PanCK+ |
| imPVL | CD146+Thy1+; CD140b+CD146+Thy1+ |
| dPVL | CD146+; CD140b+CD146; CD140b+CD146+a-SMA+ |
| Endothelial cells | CD31+; CD31+CD146+; CD31+CD140b+ |
| iCAF | CD140b+ |
| PD1-CD8+ T cells | CD8+; CD8+CD140b+; CD8+PanCK+ |
| PD1+CD8+ T cells | PD1+CD8+; PD1+CD8+CD140b+; PD1+CD8+PanCK+ |

**Supplementary table 3.** Description of features selected for assessing the association with clinicopathological variables.

| **Features** | **Description** |
| --- | --- |
| % in stroma | Proportions of CAF subsets among all detected cells in the stromal region |
| Epithelial_NMS | Normalized mixing score between CAF subsets and epithelial cells |
| CD8_NMS | Normalized mixing score between CAF subsets and CD8 T cells |
| Endothelial_NMS | Normalized mixing score between CAF subsets and endothelial cells |
| Epithelial Min Dist | Averaged distance from CAF subsets to the nearest epithelial cell |
| CD8 Min Dist | Averaged distance from CAF subsets to the nearest CD8 T cells |
| Endothelial Min Dist | Averaged distance from CAF subsets to the nearest endothelial cells |
| adj. to Epithelial | Proportions of CAF subsets located within 30µm radius to epithelial cells |
| adj. to CD8 | Proportions of CAF subsets located within 30µm radius to CD8 T cells |
| adj. to Endothelial | Proportions of CAF subsets located within 30µm radius to endothelial cells |

**Supplementary table 4.** Xenium gene list for custom-designed add-ons

| AC244453.1 | ACKR1 | ACTA2 | ALDH1A1 | ANKRD22 | ATP8B1 | C5AR2 | CCL21 | CD69 | CD74 |
| --- | --- | --- | --- | --- | --- | --- | --- | --- | --- |
| CDH19 | CHI3L1 | CISH | CLEC4E | CNN1 | COL15A1 | CPA3 | CUEDC1 | CXCL14 | CXCL8 |
| CYB561 | DAPK2 | DCN | DOCK1 | DPM2 | EBAG9 | EGLN2 | ELF3 | ELL3 | EPS15L1 |
| FBLN1 | FGF18 | FKBP4 | FKBP5 | FLOT1 | FN1 | GFPT2 | GNB1 | GREB1 | GRIP1 |
| HDAC11 | HHIP | HLA-DRA | HLA-DRB1 | HPN | HSPB1 | IGFBP4 | IGFL2 | IRF9 | ITGA10 |
| KDM4C | KLF10 | KLF2 | KRT15 | KRT19 | KRT5 | KRT8 | LGI4 | LUM | LYVE1 |
| LYZ | MEF2B | MEG8 | MEOX2 | MMP1 | MMP2 | MNDA | MTRNR2L12 | MYH11 | NID2 |
| NTAN1 | OGN | OSMR | P4HA2 | PARD6G | PARP9 | PCOLCE | PLA2G2A | PLP1 | PTPRC |
| PTPRD | RAMP3 | RASGEF1B | RGS13 | S100A8 | S100A9 | SCD | SFRP2 | SGK3 | SLC14A1 |
| SPDEF | SPNS2 | SPP1 | SRGN | SRSF5 | TAGLN | TPSB2 | UBE2B | VIM | VWF |

**Supplementary table 5.** Marker gene combinations for defining cell types in Xenium. Alternative combinations were separated by semi comma.

| **Cell type** | **Phenotype marker combinations** |
| --- | --- |
| myCAF | *ACTA2*+*PDGFRB*+ |
| Epithelial cells | *EPCAM*+; *PECAM1*+*EPCAM*+ |
| imPVL | *MCAM*+*THY1*+; *MCAM*+*THY1*+*PDGFRB*+ |
| dPVL | *MCAM*+; *PDGFRB*+*MCAM*+; *PDGFRB*+*MCAM*+*ACTA2*+ |
| Endothelial cells | *PECAM1*+; *PECAM1*+*MCAM*+; *PECAM1*+*PDGFRB*+ |
| iCAF | *PDGFRB*+ |
| PD1-CD8+ T cells | *CD8A*+; *CD8A*+*PDGFRB*; *CD8A*+*EPCAM*+ |
| PD1+CD8+ T cells | *PDCD1*+*CD8A*+; *PDCD1*+*CD8A*+*PDGFRB*+; *PDCD*+*CD8A*+*EPCAM*+ |

**Supplementary table 6:** Univariate and multivariate Cox regression of clinicopathological predictors of overall survival in in the luminal cohort. p<0.05 was considered statistically significant (HR: hazard ratio; CI: confidence interval).

|  | **Univariate analysis** | | | **Multivariate analysis** | | |
| --- | --- | --- | --- | --- | --- | --- |
|  | HR | 95% CI | P-value | HR | 95% CI | P-value |
| Age (>=55) | 3.12 | 1.95-4.97 | **<0.001** | 3.55 | 2.21-5.71 | **<0.001** |
| Node (N^+^) | 1.55 | 1.09-2.19 | **0.014** | 1.40 | 0.97-2.02 | 0.072 |
| Grade (2-Medium) | 1.58 | 1.06-2.35 | **0.023** | 1.35 | 0.89-2.04 | 0.16 |
| Grade (3-High) | 1.35 | 0.84-2.18 | 0.217 |  |  |  |
| Chemo (No) | 1.32 | 0.79-2.19 | 0.289 |  |  |  |
| Boost (No) | 1.36 | 0.96-1.91 | 0.080 |  |  |  |
| Tumour size (>=20mm) | 1.46 | 1.03-2.06 | **0.032** | 1.41 | 0.97-2.05 | 0.072 |
| Molecular subtype (luminal B) | 1.86 | 1.30-2.66 | **<0.001** | 2.07 | 1.38-3.12 | **<0.001** |

**Supplementary table 7.** Univariate and multivariate Cox regression of clinicopathological predictors of overall survival in in the TNBC cohort. p<0.05 was considered statistically significant (HR: hazard ratio; CI: confidence interval).

|  | **Univariate analysis** | | | **Multivariate analysis** | | |
| --- | --- | --- | --- | --- | --- | --- |
|  | HR | 95% CI | P-value | HR | 95% CI | P-value |
| Age (>=55) | 2.77 | 1.59-4.84 | **<0.001** | 2.75 | 1.48-5.10 | **0.0014** |
| Node (N^+^) | 2.39 | 1.47-3.91 | **<0.001** | 2.80 | 1.56-5.02 | **<0.001** |
| Chemo (no) | 2.07 | 1.23-3.47 | **0.006** | 1.86 | 1.03-3.36 | **0.039** |
| Tumour size (>=20mm) | 2.03 | 1.17-3.54 | **0.012** | 1.66 | 0.92-3.01 | 0.092 |
| TILs30 (>=30) | 0.58 | 0.35-0.97 | **0.037** | 0.56 | 0.33-0.96 | **0.036** |

**Supplementary Figure 1.** Association of CD8 T cell percentages with clinical characteristics in the luminal cohort. Clinical variables include age at diagnosis (**A**), node metastasis (**B**), tumour grade (**C**), molecular subtypes (**D**), and tumour size (**E**).

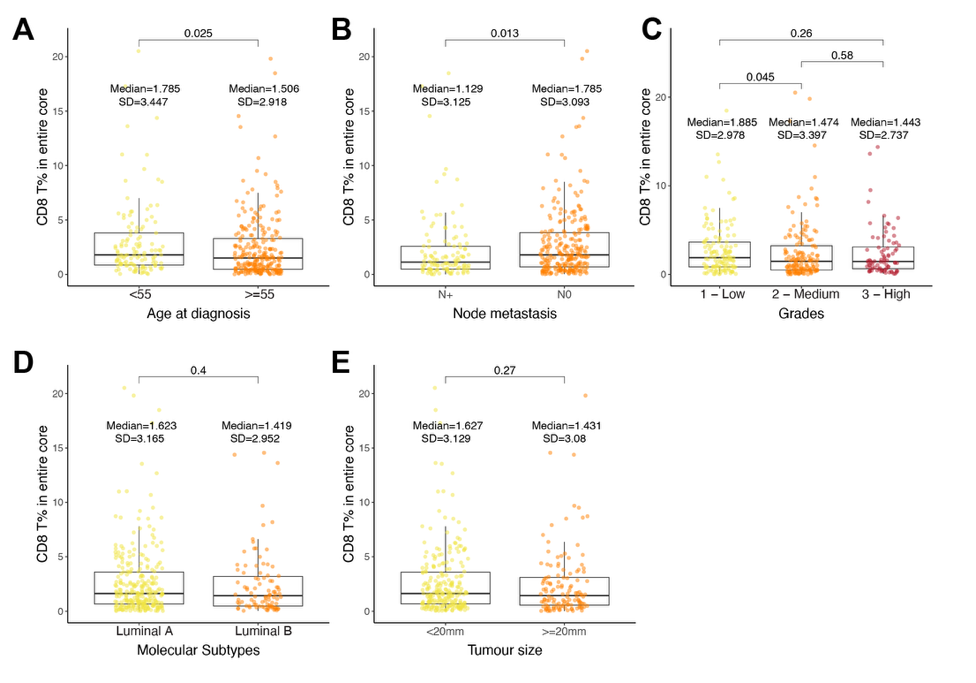

**Supplementary Figure 2. A.** Percentage of PVLs with endothelial cells present within a 10–100 µm radius in the TNBC cohort. **B, C**. Correlation between the percentage of distal endothelial cells and distal PVLs in luminal (**B**) and TNBC (**C**) cohorts. Disseminated endothelial cells are defined as those without a PVL within 100 µm.

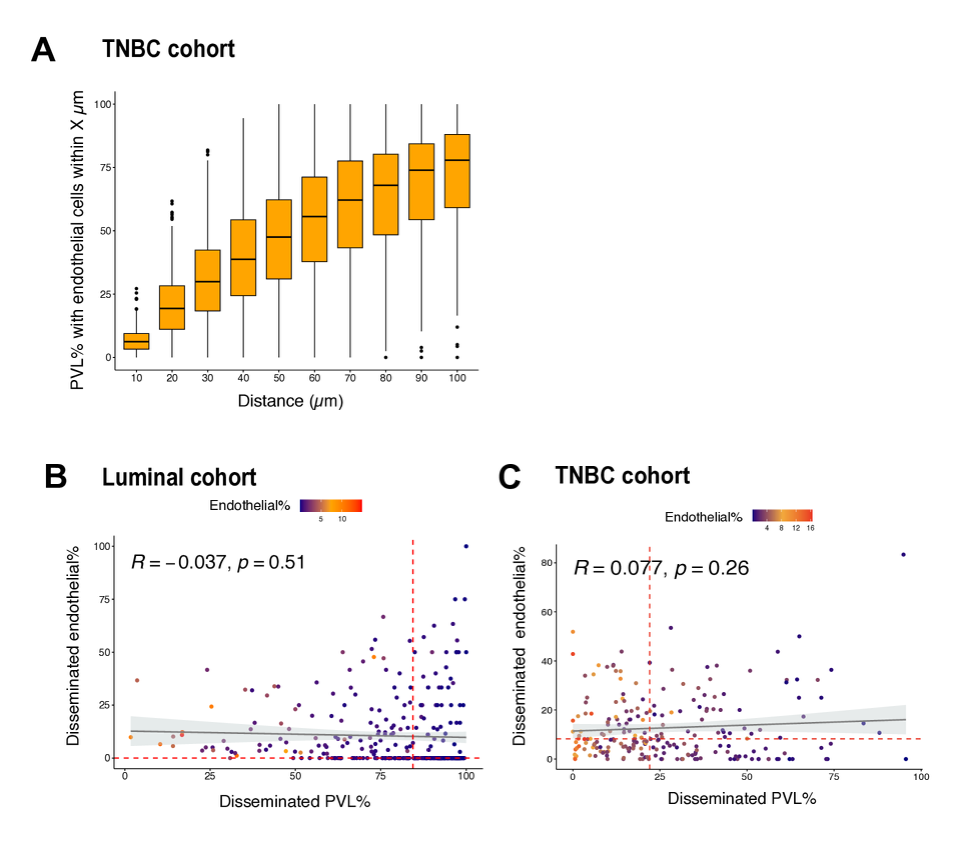

**Supplementary Figure 3.** **A.** KM curves for OS stratified by the percentage of PD1⁻CD8⁺ T cells in myCAF-high vs. myCAF-low groups. **B.** Multivariable Cox proportional hazard regression analysis of OS for the proportion of PD1⁻CD8⁺ T cells% in myCAF-low group, considering age, lymph node metastasis, chemotherapy and tumour size.

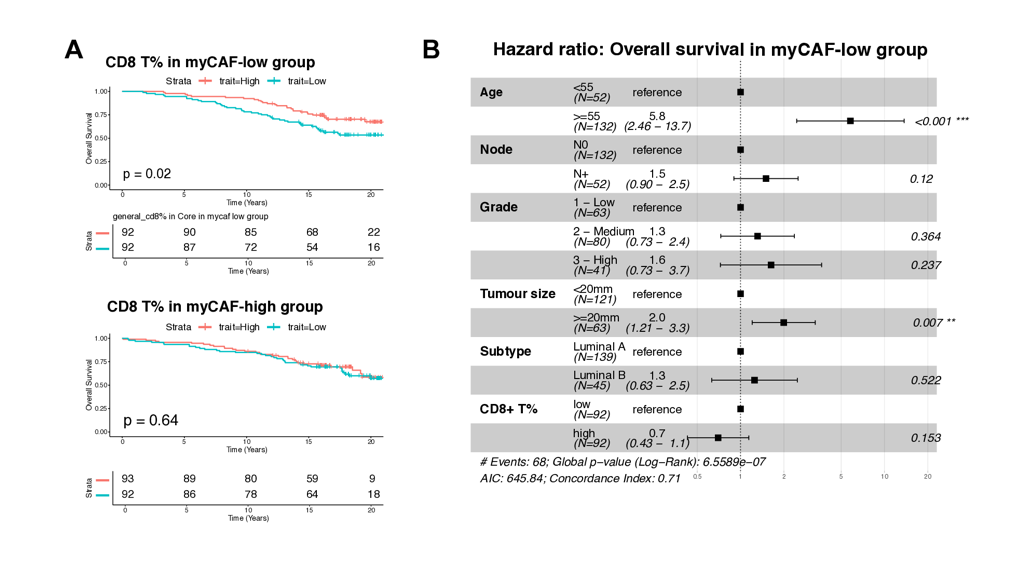

**Supplementary Figure 4. A.** Association of myCAF-related spatial features with clinical variables in the luminal cohort. Patients were stratified by clinical variables. Dot colors indicate patient groups in which the feature value is increased. Dot outlines denote statistical significance of feature enrichment (red: significant; grey: not significant), assessed by the Wilcoxon rank-sum test with p-values corrected using the Benjamini-Hochberg method. **B.** Endothelial cell percentages in the stromal region across tumours of 1 to 3 grades. **C.** Association of myCAF-related spatial features with clinical variables in the TNBC cohort.

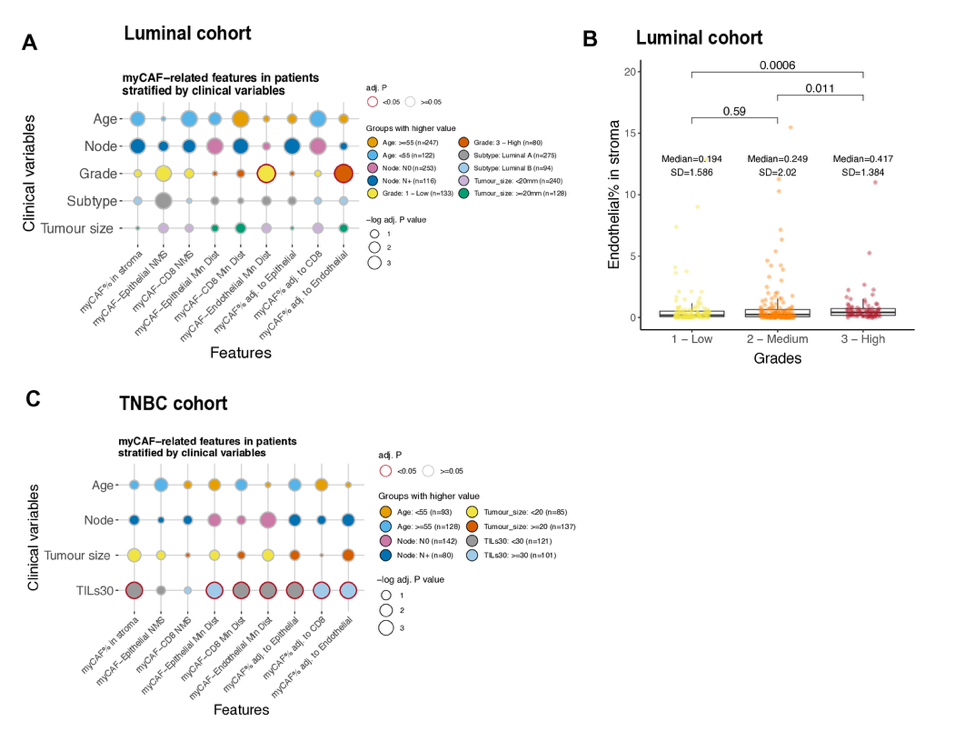
